# SARS-CoV-2 replication triggers an MDA-5-dependent interferon production which is unable to efficiently control replication

**DOI:** 10.1101/2020.10.28.358945

**Authors:** Rebendenne Antoine, Chaves Valadão Ana Luiza, Tauziet Marine, Maarifi Ghizlane, Bonaventure Boris, Planès Rémi, McKellar Joe, Nisole Sébastien, Arnaud-Arnould Mary, Moncorgé Olivier, Goujon Caroline

## Abstract

Severe acute respiratory syndrome coronavirus 2 (SARS-CoV-2) is the third highly pathogenic coronavirus to spill over to humans in less than 20 years, after SARS-CoV-1 in 2002-2003 and Middle East respiratory syndrome (MERS)-CoV in 2012. SARS-CoV-2 is the etiologic agent of coronavirus disease 19 (COVID-19), which ranges from mild respiratory symptoms to severe lung injury and death in the most severe cases. The COVID-19 pandemic is currently a major health issue worldwide. Immune dysregulation characterized by altered innate cytokine responses is thought to contribute to the pathology of COVID-19 patients, which is a testimony of the fundamental role of the innate immune response against SARS-CoV-2. Here, we further characterized the host cell antiviral response against SARS-CoV-2 by using primary human airway epithelia and immortalized model cell lines. We mainly focused on the type I and III interferon (IFN) responses, which lead to the establishment of an antiviral state through the expression of IFN-stimulated genes (ISGs). Our results demonstrate that both primary airway epithelial cells and model cell lines elicit a robust immune response characterized by a strong induction of type I and III IFN through the detection of viral pathogen molecular patterns (PAMPs) by melanoma differentiation associated gene (MDA)-5. However, despite the high levels of type I and III IFNs produced in response to SARS-CoV-2 infection, the IFN response was unable to control viral replication, whereas IFN pre-treatment strongly inhibited viral replication and *de novo* production of infectious virions. Taken together, these results highlight the complex and ambiguous interplay between viral replication and the timing of IFN responses.

---

Severe acute respiratory syndrome coronavirus 2 (SARS-CoV-2) is responsible for the current coronavirus disease 2019 (COVID-19) pandemic. This virus emerged in China at the end of 2019 (Zhou et al., 2020a; Zhu et al., 2020) and has, since then, dramatically spread across the world. As of today (October 28th, 2020), SARS-CoV-2 has caused more than 1,172,000 deaths worldwide, general lockdowns in many countries across the world and an unprecedented global economic crisis, as well as an undeniable pressure on health systems. SARS-CoV-2 is closely related to the highly pathogenic SARS-CoV-1, which caused an outbreak in 2002-2003 in South East China and Hong Kong, with a fatality rate close to ~10% (Drosten et al., 2003; Peiris et al., 2003; Zhong et al., 2003). COVID-19 has many of the hallmarks of SARS-CoV disease including fever, breathing difficulty, acute respiratory distress syndrome (ARDS) and death in the most severe cases (Huang et al., 2020). SARS-CoV-2 is also related to Middle East Respiratory Syndrome coronavirus (MERS-CoV), another highly pathogenic coronavirus discovered in 2002, causing an ARDS and organ failure in the most severe cases, and death in ~35% of cases (Zaki et al., 2012). The estimated fatality rate of SARS-CoV-2 infection is much lower, but still fairly high (estimated to be ~0.5-1%), and this virus has clearly shown a great potential for transmission between humans. Four additional coronaviruses, known to infect humans, circulate in the population every winter, mainly causing common colds: the human coronaviruses (HCoV)-229E, -OC43, -NL63, and -HKU1.

Together with SARS-CoV-1 and MERS-CoV, SARS-CoV-2 belongs to the genus *betacoronavirus*. They are enveloped viruses with large positive-sense RNA genomes of approximately 30 kilobases. The genome of SARS-CoV-2 shares 79.6% identity with SARS-CoV-1 and 50% with MERS-CoV (Zhou et al., 2020a). SARS-CoV-2 mainly replicates in the respiratory tract, but can also replicate in the gastrointestinal tract (Xiao et al., 2020). Similarly to SARS-CoV-1 and HCoV-NL63, SARS-CoV-2 entry into target cells is mediated by the Angiotensin converting enzyme 2 (ACE2) receptor (Hoffmann et al., 2020; Hofmann et al., 2005; Li et al., 2003; Zhou et al., 2020b). Recently, Neuropilin-1 was identified as an important cofactor implicated in SARS-CoV-2 entry (Cantuti-Castelvetri et al., 2020). The cellular serine protease Transmembrane protease, serine 2 (TMPRSS2) is employed by both SARS-CoV-1 and −2 for Spike (S) protein priming at the plasma membrane (Hoffmann et al., 2020; Matsuyama et al., 2010). Cathepsins may also be involved in SARS-CoV S cleavage and fusion peptide exposure, upon entry via an endocytic route (Huang et al., 2006; Ou et al., 2020; Simmons et al., 2005).

Viral infections are detected by the innate immune system through the recognition of pathogen-associated molecular patterns (PAMPs) by pattern recognition receptors (PRRs). PAMPs, which are typical features not found in host cells, are present in the pathogen or generated during infection, and are recognized by at least three distinct classes of PRRs: the Toll-like receptors (TLR), the Retinoic acid-inducible gene I (RIG-I)-like receptors RIG-I and melanoma differentiation associated gene (MDA)-5, and the nucleotide-binding oligomerization domain-like receptors (NOD)-like receptor family member (NLRs), such as NLR family pyrin domain containing 3 (NLRP3). These PRRs share different subcellular localizations, with NLRP3 and RIG-I/MDA-5 located in the cytoplasm, and TLR present within endosomes or at the plasma membrane (Park and Iwasaki, 2020). Genetic studies have shown that mouse hepatitis virus (MHV), a murine coronavirus, is sensed by both RIG-I and MDA-5, with a critical role of the latter *in vivo* (Li et al., 2010; Park and Iwasaki, 2020; Sa Ribero et al., 2020; Zalinger et al., 2015). TLR3 seems to play an important role in the sensing of MHV, SARS-CoV-1 and MERS-CoV in plasmacytoid dendritic cells (Cervantes-Barragan et al., 2007, Scheuplein et al., 2015). However, the role of known PRRs in SARS-CoV-2 sensing has yet to be elucidated.

PRR activation leads to the production and secretion of type I and III interferons, pro-inflammatory cytokines, eicosanoids and chemokines, which can act in a paracrine and autocrine manner. In particular, type I and III IFNs constitute one of the first lines of defence against viral infections, through the regulation of hundreds of interferon-stimulated genes (ISGs), which induce an antiviral state in infected and neighbouring cells. Type I IFNs include 17 subtypes (13 IFN-α, IFN-β, IFN-ε, IFN-κ, IFN-ω) and type III IFNs include four subtypes: IFN-λ1 (IL-29), IFN-λ2 (IL-28A), IFN-λ3 (IL-28B), and IFN-λ4 (Lazear et al., 2019). Receptors for type I IFN (IFNAR1 and 2) are ubiquitously expressed, whereas IFN-III receptors are preferentially expressed on mucosal epithelial cells, as well as liver cells and some myeloid cells (Kotenko et al., 2019; Wack et al., 2015). Both type I and III IFNs induce ISGs but type I IFN leads to a more rapid induction and decline of ISG expression and is more pro-inflammatory than type III (Lazear et al., 2019).

Type I and III IFNs treatments have been shown to inhibit the replication of SARS-CoV-1 (Haagmans et al., 2004; Paragas et al., 2005; Zheng et al., 2004), and more dramatically that of MERS-CoV (Chan et al., 2015; Hart et al., 2014; de Wilde et al., 2013) and SARS-CoV-2 (Blanco-Melo et al., 2020; Dinnon et al., 2020; Hassan et al., 2020; Israelow et al., 2020a; Lei et al., 2020a; Lokugamage et al., 2020; Stanifer et al., 2020), both *in vitro* and *in vivo*. However, SARS-CoV-2 dampens type I and III IFN induction and subsequent expression of ISGs (Blanco-Melo et al., 2020; Israelow et al., 2020b; Lei et al., 2020a, 2020b). Upon SARS-CoV-1 infection of mice, a similar phenotype was observed and was associated with a delayed type I IFN signalling, which was linked to disease severity. Indeed, type I IFN promoted accumulation of pathogenic monocyte-macrophages, resulting in lung immunopathology, vascular leakage, and suboptimal T cell responses, whereas early type I IFN administration or impairment of type I IFN signalling at later stages upon infection ameliorated immunopathology (Channappanavar et al., 2016). Consistent with this, the recruitment of proinflammatory cells was dependent on type I IFN signalling, but not the viral clearance in a murine model of infection by SARS-CoV-2 (Israelow et al., 2020a), suggesting that IFNs might be more deleterious than beneficial *in vivo*. However, in COVID-19 patients, the levels of circulating IFN-β were undetectable and an impaired IFN-α production seemed to characterize the most severe cases (Arunachalam et al., 2020; Hadjadj et al., 2020). Moreover, neutralizing auto-antibodies against type I IFNs were found in at least 10% of life-threatening COVID-19 patients (Bastard et al., 2020). Furthermore, 3,5% of the critically ill patients with severe COVID-19 carried inborn errors in genes involved in the type I IFN pathway such as *TLR3*, *IRF7* or *IFNAR1* (Zhang et al., 2020). Taken together, this highlights a critical role of IFNs in COVID-19 disease severity and the importance of better understanding the interplay between SARS-CoV-2 and the IFN system.

In this article, we aimed to characterize cell host responses to SARS-CoV-2 infection and the role of IFNs in the control of viral replication, using primary human airway epithelia and immortalized model cell lines. First, we showed that primary human airway epithelial cells supported high levels of SARS-CoV-2 replication and were capable of sensing this virus, with a potent induction of genes involved in innate immunity, including type I and III IFNs, and production of these cytokines in their basal media. However, this response arrived late upon viral exposure (after 48 h). The naturally permissive Calu-3 lung cell line recapitulated IFN induction upon SARS-CoV-2 exposure, which validated the use of this model cell line as a tool to study SARS-CoV-2 replication and induction of innate immunity. ACE2-transduced, lung A549 and intestinal Caco-2 cell lines were also able to detect SARS-CoV-2 virus, albeit with different efficiencies. Interestingly, all these cell types could inhibit SARS-CoV-2 replication to different levels upon type I IFN pre-exposure. Using Calu-3 cells, we further showed that MDA-5 was the main innate immune sensor of SARS-CoV-2 in these epithelial cells. Finally, we demonstrated that type I and III IFN production elicited by SARS-CoV-2 infection was unable to inhibit replication, supporting the idea that the timing of IFN exposure is key to control replication.

## Results

### SARS-CoV-2 replicates efficiently in primary HAE cells while triggering type I and type III IFN responses

SARS-CoV-2 BetaCoV_France_IDF0372_2020_C2 isolate (obtained from Pasteur Institute) was amplified and titrated on Vero E6 cells by plaque assays. In order to analyse host cell responses to SARS-CoV-2 infection in physiological targets, 3D Human Airway Epithelia (HAE; MucilAir™ cells, Epithelix) were mock-infected or incubated with SARS-CoV-2 on the apical side at a MOI of 0,01 or 0,1. Viral replication was analysed at the indicated time points, by monitoring the copy number of RNA polymerase RNA dependent (RdRp) RNAs in cells by RT-qPCR (Figures 1A and S1A). As shown previously (Pizzorno et al., 2020), these primary cells were highly permissive to SARS-CoV-2 replication. Immunofluorescence analyses confirmed detection of double-stranded RNA in SARS-CoV-2 infected HAE cells but not in non-infected cells (Figure 1B). The basal media was harvested at 72 h post-infection, and cytokine production was measured using the Human Anti-Virus Response Panel LEGENDplex™ (Figure 1C). A strong IFN response was observed, with a massive production of both type I (IFN-β) and type III (IFN-γ1, and 2/3) IFNs. An important induction of CXCL10 (also named IP-10) production was also observed, along with a more modest induction of pro-inflammatory cytokines IL-6, IL-8 and TNF-α. A RT-qPCR experiment on samples collected at 24, 48 and 72 h post-infection confirmed an important induction of *IFNB1, IFNL1 and L2* at 48 h and 72 h after infection with SARS-CoV-2, which was not observed at 24 h, and parallel induction of prototype ISGs (Figure S1B). Next, a RT^2^ profiler analysis using the Antiviral Response panel was performed at 72 h post infection (Figure 1D). As expected, *IFNB* and other genes belonging to the IFN system (e.g. *STAT1*, *TICAM1*, *TLR3*, *TLR7*, *TLR8*, *DDX58*, *IRF7*, as well as antiviral effector ISGs, such as *OAS2* and *MX1*) were confirmed to be upregulated.

**Figure 1.**
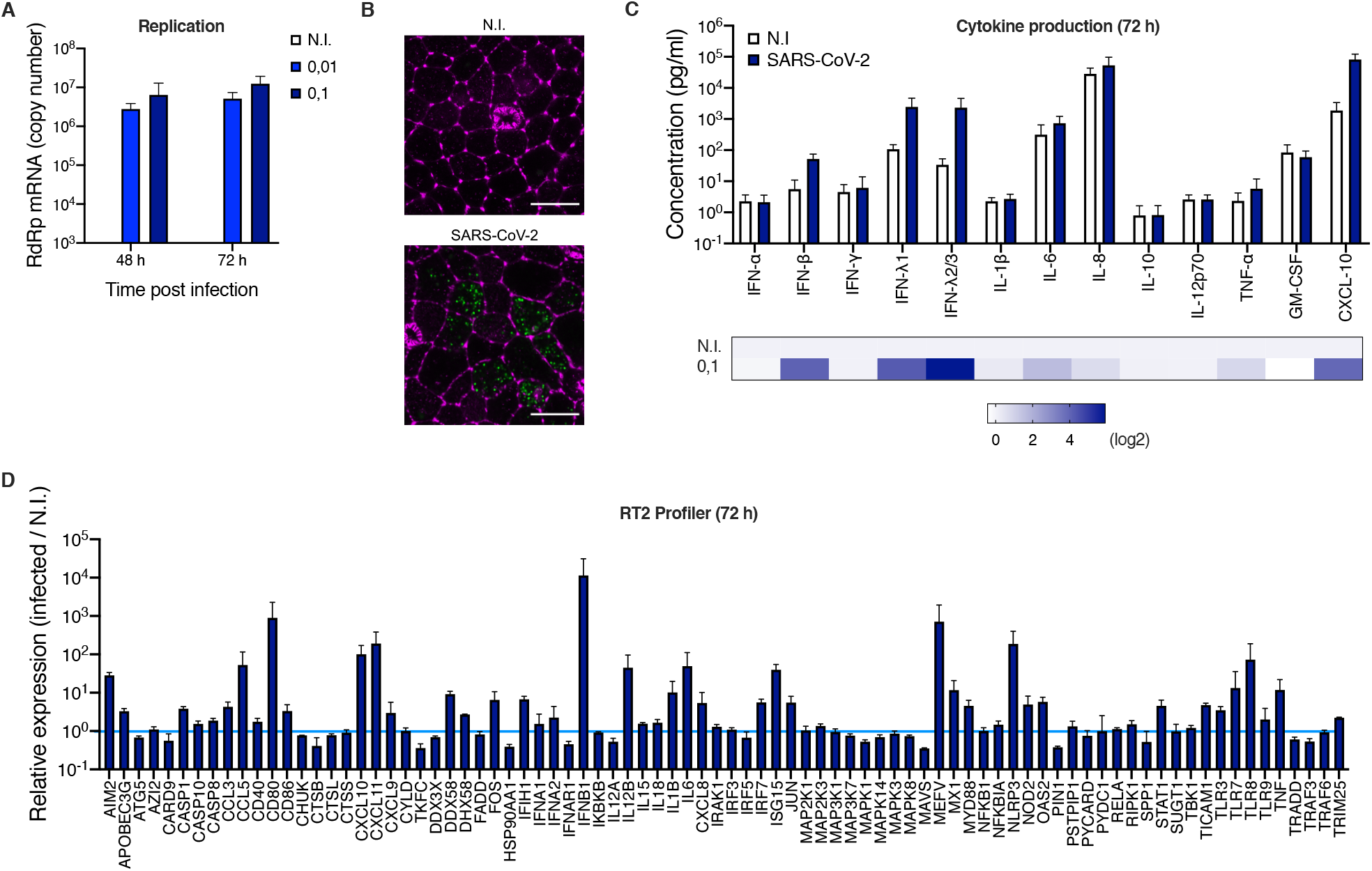
Primary human airway epithelial host cell responses to SARS-CoV-2 infection. **A**. Human HAE cells (MucilAir™, Epithelix) were mock infected (N.I.) or incubated with SARS-CoV-2 on the apical side at MOI 0,01 and 0,1 for 2 h. Viral input was removed and the apical side washed with PBS1X before incubation for 48 h and 72 h. Cells were harvested at the indicated time points and lysed for RNA extraction and RT-qPCR analysis using RdRp primers and probe. **B**. Human HAE cells were mock infected (N.I.) or incubated with SARS-CoV-2 on the apical side at MOI 0,1 for 2 h. Viral input was removed and the apical side washed with PBS1X before incubation for 48 h. Cells were fixed and stained for Actin with Phalloidin conjugated to Alexa Fluor 488 (shown here in magenta) and with an anti-double stranded RNA and a secondary antibody conjugated to Alexa Fluor 546 (shown in green). Images were acquired with an LSM880 Airyscan microscope. Representative images are shown; scale bar 10 μm. **C**. Human HAE cells were mock infected (N.I.) or incubated with SARS-COV-2 (MOI 0,1), as in A. Cytokine concentrations in the basal media were measured using the Human Anti-Virus Response Panel LEGENDplex™ at 72 h post-infection (top) and the fold difference in cytokine concentration in the basal media from infected compared to N.I. cells is represented as a heat map (bottom; log2 scale). **D**. An Antiviral Response RT^2^ profiler PCR array analysis was performed using the RNAs extracted at 72 h (MOI 0,1). The mean of 4 (A) or 3 (C and D) independent experiments is shown, with error bars representing one standard deviation (s.d.) from the mean.

A potent induction of the inflammasome genes *NLRP3* and *MEFV* (encoding Pyrin, also known as TRIM20) was interestingly also observed, as well as CXCL10, CXCL11, CCL3 and CCL5 chemokine induction. Of note, HAE cells from nasal, tracheal and bronchial origins behaved globally similarly with respect to their responses to infection (Figure S1C and D).

### Calu-3 cells recapitulate the global responses to SARS-CoV-2 observed in HAE cells

Next, a number of immortalized, epithelial human cell lines were tested for their ability to support viral replication, in comparison to simian Vero E6 cells (Figure S2). Calu-3 (lung adenocarcinoma), Caco-2 (colorectal adenocarcinoma) (both known to express SARS-CoV-2 receptor ACE2 and TMPRSS2 protease and be permissive to SARS-CoV-2 (Chu et al., 2020) were used in parallel to A549 (epithelial, lung carcinoma) genetically modified to express ACE2, together with TMPRSS2 or not. We also genetically modified Caco-2 cells to express higher levels of ACE2, or ACE2 and TMPRSS2 in combination. The cells were infected with SARS-CoV-2 and lysed 48 h later to measure viral replication using a RdRp RT-qPCR. Calu-3, Caco-2-ACE2 and A549-ACE2 supported SARS-CoV-2 replication to a very similar extent (Figure S2) and were therefore selected for further studies.

Responses to infection were next evaluated in Calu-3 cells (Figure 2). The cells were infected at the indicated MOIs and replication efficiency was assessed by measuring RdRp copy numbers (Figure 2A). Whereas replication efficiency increased concomitantly with the viral input at 24 h, a plateau was reached at 48 h. Cytokine production in Calu-3 supernatants was measured and a very similar response than that in HAE cells was observed, with a high induction of IFN-β, IFN-γ1 and IFN-γ2/3, CXCL10 and a slight induction of IL-6 and TNF-α(Figure 2B). RT^2^ profiler and RT-qPCR analyses confirmed a globally similar response of Calu-3 cells to SARS-CoV-2 infection as compared to HAE cells (Figure 2C and Figure S3A), with a high induction of *IFNB1*, *IFNL2* and a moderate induction of ISGs. However, these responses happened more rapidly than in HAE cells (Figure S1B and S3A). Type I and III IFN production in supernatants from infected Calu-3 cells was confirmed using reporter cell lines (Figure S3B and C). High and moderate induction levels of *IFNB1*, *IFNL2*, and prototype ISGs *ISG15* and *MX1* were observed in infected A549-ACE2 and Caco2-ACE2, respectively, upon SARS-CoV-2 replication (Figure S4).

**Figure 2.**
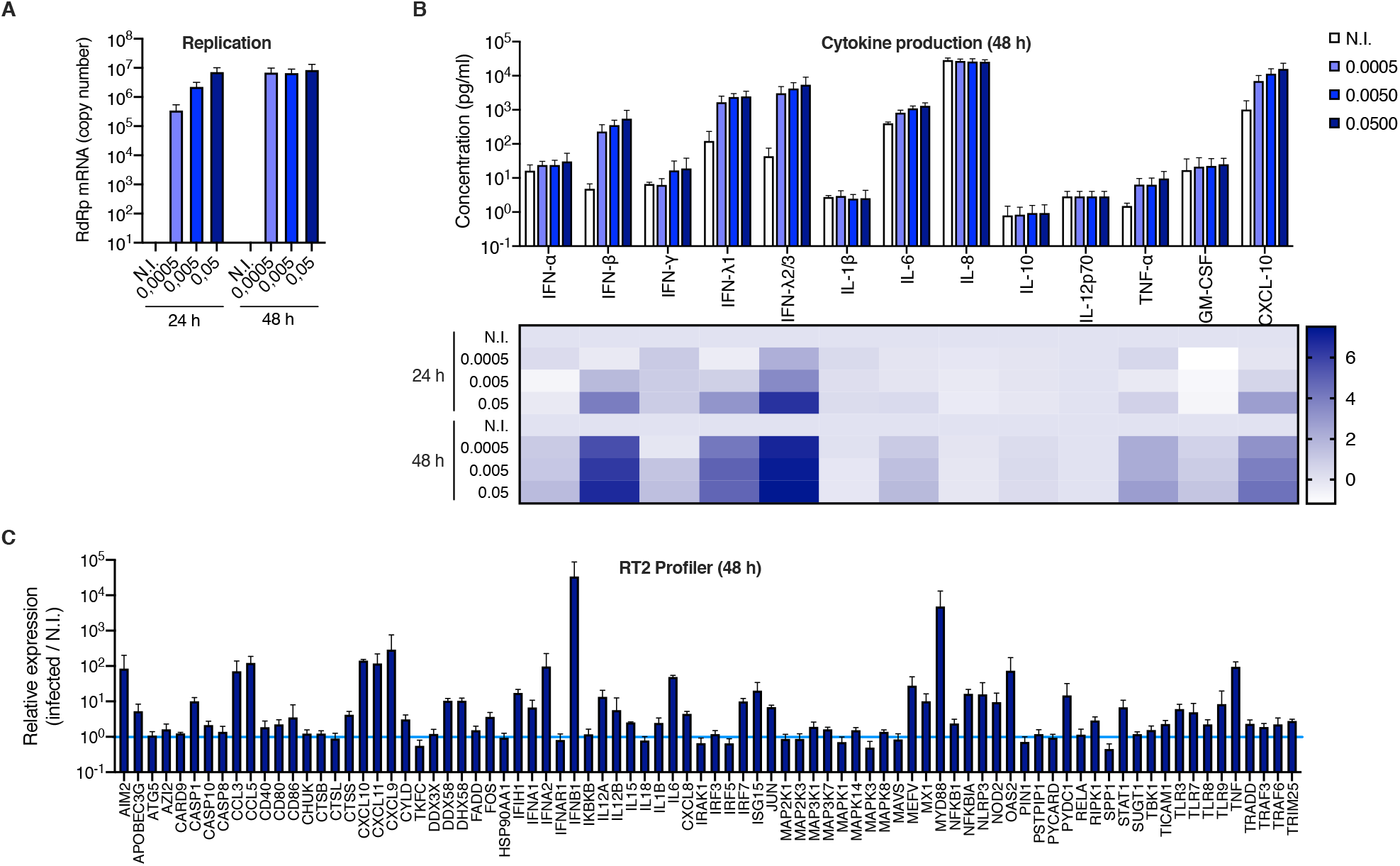
Calu-3 model cell line responses to SARS-CoV-2 infection. **A**. Human Calu-3 cells were non-infected (N.I.) or incubated with SARS-CoV-2 at MOIs of 0,0005, 0,005 and 0,05. Cells were harvested at the indicated time points and lysed for RNA extraction and RT-qPCR analysis using RdRp primers and probe. **B**. The cell supernatants from (A) were harvested at the indicated time points and cytokine concentrations were measured using the Human Anti-Virus Response Panel LEGENDplex™ at 24 h and 48 h. Concentrations are shown (top), and the fold difference in cytokine concentration in supernatants from infected compared to non-infected cells was calculated and represented as a heat map at 24 h and 48 h (bottom; log2 scale). **C**. An Antiviral Response RT^2^ profiler PCR array analysis was performed using the RNAs extracted at 48 h (MOI 0,005). The mean of 4 (A) or 3 (B and C) independent experiments is shown, with error bars representing one s.d. from the mean.

Having established that SARS-CoV-2 infected cells produced high amounts of IFN, we sought to identify the PRR(s) responsible for sensing the virus. Typically, coronaviruses are sensed via RIG-I and/or MDA-5 PRRs (Li et al., 2010; Zalinger et al., 2015), which then signal through mitochondrial antiviral-signalling protein (MAVS). We therefore used CRISPR/Cas9 to generate cell populations knockout for RIG-I, MDA-5 or MAVS and tested the ability of these cells to produce type I and III IFNs upon SARS-CoV-2 infection (Figure 3). We observed that MDA-5 and MAVS depletion, but not RIG-I depletion, drastically impacted the amounts of type I and type III IFNs produced, demonstrating that in Calu-3 lung epithelial cells, SARS-CoV-2 sensing mainly occurred through MDA-5.

**Figure 3.**
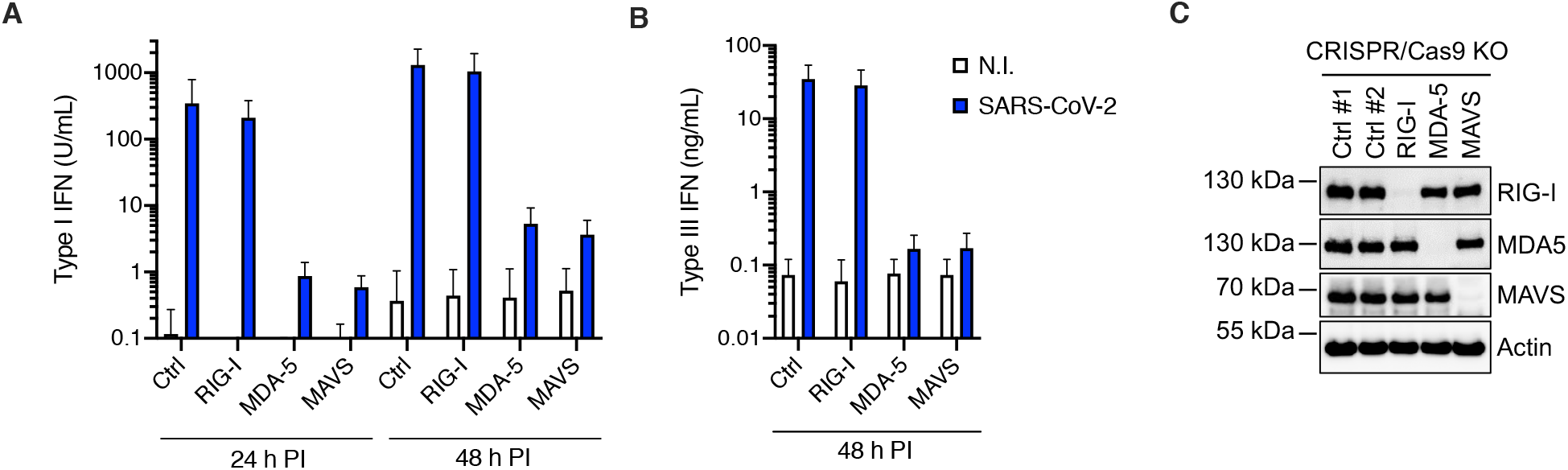
MDA-5 is the main sensor of SARS-CoV-2 in Calu-3 model cells. Calu-3-Cas9 cells were transduced with lentiviral vectors expressing CRISPR non-targeting single guide RNAs (Ctrl #1, #2) or single guide RNAs targeting RIG-I, MDA-5 or MAVS. Cells were antibiotic selected for at least 15 days and challenged with SARS-CoV-2 at MOI 0,05. The cell supernatants were harvested at 24 h and 48 h post-infection, as indicated, and the concentrations of type I (**A**) and type III (**B**) IFNs produced analysed using HEK-Blue™ IFN-α/β and IFN-γ reporter cells, respectively. The mean of 3 independent experiments (performed with 2 series of KO cell populations generated independently) is shown, with error bars representing one s.d. from the mean. **C**. A representative immunoblot is shown, Actin served as a loading control.

In agreement with previous studies (Felgenhauer et al., 2020; Lokugamage et al., 2020; Mantlo et al., 2020), we then observed that a 24 h pre-treatment with increasing doses of type I IFN proportionally limited SARS-CoV-2 replication in Vero E6 cells, with the best dose being 1000 U/mL (Figure S5). Interestingly, pre-exposure of HAE and Calu-3 cells with 1000 U/mL IFN potently decreased SARS-CoV-2 RNA amounts in infected cells (by 1,5-2 logs) and the production of infectious viruses (by several orders of magnitude; Figure 4A-B and D-E). Immunofluorescence staining failed to detect dsRNA in SARS-CoV-2 infected cells following IFN pre-treatment in HAE cells, contrary to what was observed in non-treated cells (Figure 4C). Intracellular spike staining in infected Calu-3 cells showed a 10-fold decrease in the percentage of infected cells following IFN pre-exposure (Figure 4F) and immunoblot analysis confirmed a potent inhibition of Nucleocapsid (N) and Spike expression, concomitant with ISG induction (Figure 4G). Of note, type I IFN pre-treatment had also a very strong impact on infection in A549-ACE2 cells but a milder effect in Caco-2-ACE2 cells (Figure S6). It was quite striking that the concentrations of type I IFN used here for pre-treatment were similar to what was naturally produced by the infected cells (Figures 1C, 2A and S3A), however high levels of replication were observed in the absence of exogenous IFN treatment. We therefore hypothesized that the IFN produced during the course of infection did not have an impact on replication. Indeed, we observed that MDA-5 and MAVS knockout in Calu-3 cells did not impact viral production, despite preventing IFN production (Figure S7 and Figure 3). In order to confirm this, we used CRISPR/Cas9 to disrupt genes belonging to the common signalling pathway for type I and III IFNs, rather than the distinct type I and III IFN receptors. Hence, we generated IRF9 and control (CTRL) KO Calu-3 cell populations and we observed that IRF9 knockout did not substantially improved SARS-CoV-2 replication (Figure 4H), whereas the KO cells were not able anymore to induce prototype ISG expression following IFN treatment (Figure 4I). Similar data were obtained in *JAK1* knockout A549-ACE2 cells and similarly unable to respond to IFN (Figure S8). In line with these results, we observed that contrary to pre-exposure, IFN exposure 24 h post-infection did not have an impact on SARS-CoV-2 replication efficiency (Figure S9). Moreover, when added as early as 8 h post infection, IFN treatment had an impact only at the lowest MOIs used (Figure S9).

**Figure 4.**
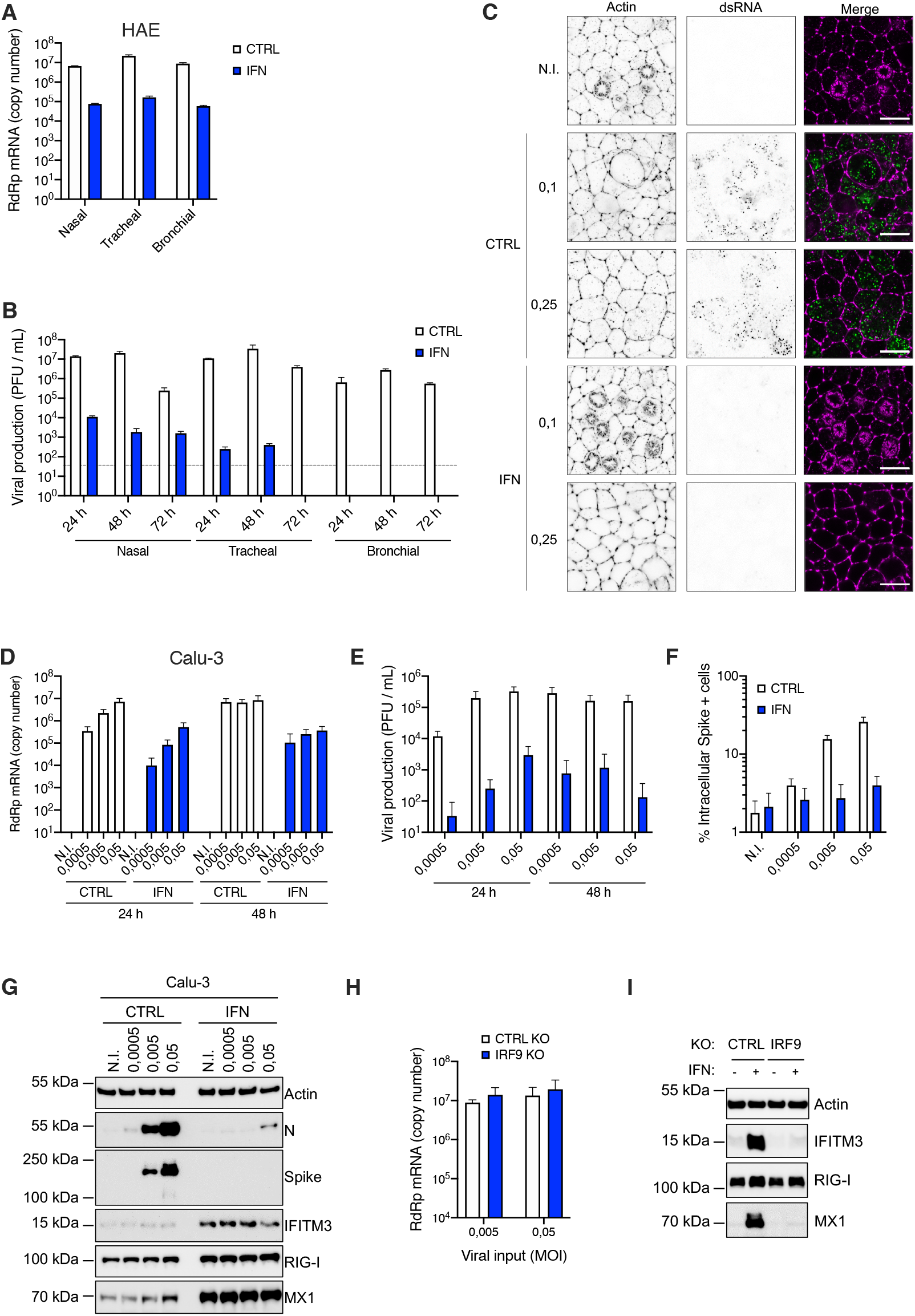
Inhibition of SARS-CoV-2 replication by type I IFN pre-treatment in primary HAE cells and immortalized Calu-3 cells. **A**. Human HAE cells of nasal, tracheal and bronchial origins (as indicated) were pre-treated or not with type I IFN for 20 h, and mock infected (N.I.) or incubated with SARS-CoV-2 on the apical side at MOI 0,1 for 1-2 h. The viral input was removed and the apical side washed in PBS1X. Cells were harvested and lysed for RNA extraction and RT-qPCR analysis using RdRp primers and probe at 72 h post-infection. **B**. Washes of the apical side of the HAE cells from (A) were performed at 24 h, 48 h and 72 h, harvested and frozen down for subsequent titrations. Plaque assays were then performed in technical duplicates to determine the number of plaque forming units (PFU) per mL of supernatant in each condition. The grey, dotted line indicates the detection threshold. **C**. Human HAE cells of bronchial origin were pre-treated or not with IFN for 20 h, and mock infected (N.I.) or incubated with SARS-CoV-2 on the apical side at MOI 0,1 and 0,25, as indicated, for 2 h. Viral input was removed and the apical side washed with PBS1X before incubation for 48 h. Cells were fixed and stained for Actin with Phalloidin conjugated to Alexa 488 (shown in magenta on the merge images) and with an anti-double stranded RNA and a secondary antibody conjugated to Alexa 546 (shown in green on the merge images). Images were acquired with an LSM880 Airyscan microscope. Representative images are shown; scale bar 10 μm. **D**. Human Calu-3 cells were pre-treated or not with IFN for 16-20 h, the media was replaced and the cells were mock-infected (N.I.) or incubated with SARS-CoV-2 at the indicated MOIs. Cells were harvested at the indicated time points and lysed for RNA extraction and RT-qPCR analysis using RdRp primers and probe. **E**. Aliquots of the supernatants from D were harvested at 24 h and 48 h post infection and plaque assays were performed to evaluate the production of infectious viruses in the different conditions. **F**. Calu-3 cells were pre-treated or not with IFN and infected as in D, and the cells were fixed with PFA at 24 h post-infection, permeabilized and stained with an anti-Spike antibody conjugated to an Alexa fluorochrome. The percentage of Spike + cells was scored by flow cytometry. **G**. Calu-3 cells were pre-treated or not with IFN and infected as in D and lysed 24 h post-infection for immunoblot analysis of SARS-CoV-2 Nucleoprotein (N) and Spike, and IFITM3, RIG-I and MX1 expression levels, Actin serving as a loading control. A representative immunoblot is shown. **H**. CTRL and IRF9 Calu-3 knockout cells were infected with SARS-CoV-2 at the indicated MOIs and viral replication was measured 48 h later by RdRp RT-qPCR. **I**. CTRL and IRF9 knockout cells were pre-treated or not with IFN for 48 h, lysed and the expression levels of IFITM3, RIG-I and MX1 were analysed by immunoblot, Actin served as a loading control. A representative immunoblot is shown. The mean of 3 (A-B, H) or 4 (D-F) independent experiments is shown, with error bars representing one standard deviation (s.d.) from the mean.

## Discussion

Here, we characterized host cell responses to SARS-CoV-2 replication in primary, air-liquid HAE cultures and in model cell lines. We confirmed the potent induction of innate responses following infection of HAE cells with SARS-CoV-2 (Pizzorno et al., 2020), with an important but somewhat late induction of type I and III IFNs. In contrast to our results, a lack of IFN response in HAE cells exposed to SARS-CoV-2 was recently reported (Vanderheiden et al., 2020). Of note, viral production in the HAE model we used was several magnitudes of order higher than what was reported in the other study (up to 107 PFU/mL in washes from the apical side, Figure 4B of this study, in comparison to ~2-3.102 PFU/mL, (Vanderheiden et al., 2020)), which could explain the observed difference in sensing, in addition to inherent differences in the HAE models used. A lack of IFN induction was also reported upon SARS-CoV-2 infection of normal human bronchial epithelial (NHBE) cells (Blanco-Melo et al., 2020). However, in this study, only 0.1% of total deep sequencing reads were from the virus in this cell type, which was a percentage highly similar to what was observed in wild-type A549 cells (i.e. devoid of Ace2) (Blanco-Melo et al., 2020), known to be refractory to replication because of the lack of ACE2 expression. This strongly suggested poor replication efficiency in this particular model, again explaining the lack of sensing. In support of our data, and in addition to the aforementioned previous report in HAE cells (Pizzorno et al., 2020), an IFN induction has also been reported in human intestinal organoids (Lamers et al., 2020; Stanifer et al., 2020). Using model cell lines, we notably showed that naturally permissive, lung epithelial Calu-3 cells were a good model for innate immune responses to SARS-CoV-2 infection, with a similar pattern of innate immunity gene induction and pro-inflammatory cytokine production to what we observed in HAE cells. However, the IFN induction kinetics were faster in Calu-3 cells in comparison to primary cells, with a robust induction of IFN genes 24 h post-infection (versus 48 h in HAE cells). In addition to IFNs, SARS-CoV-2 infection induced the production of the CXCL10 chemokine and the pro-inflammatory cytokines TNF-α and IL-6, in both cell types (and, in HAE only, and to a lower extent, IL-8), but no production of IL-1β, consistent with observations in COVID-19 patient samples (Del Valle et al., 2020). Interestingly, at the RNA level, a potent induction of inflammasome-related genes (i.e. *NLRP3* and *MEFV*) was observed. It will be of high interest to further explore the potential regulation of the inflammasome by SARS-CoV-2 and determine whether it is activated and, if that were the case, why there is no IL-1β production by the infected cells.

In agreement with an important role of MDA-5 in host responses to MHV infection in mice (Zalinger et al., 2015), CRISPR/Cas9 knockout approaches showed that MDA-5 was the main sensor for SARS-CoV-2 in Calu-3 lung epithelial cells, with no impact of RIG-I in this particular model. Whether that would be the case in primary HAE cells remained to be determined, but it has so far proven difficult to genetically modify these cells.

As reported previously in cell lines and in models of primary bronchial epithelial, air-liquid cell cultures (Blanco-Melo et al., 2020; Busnadiego et al., 2020; Lei et al., 2020a; Lokugamage et al., 2020; Nchioua et al., 2020), we confirmed that type I IFN pre-treatment potently inhibited SARS-CoV-2 replication in primary HAE cells and in lung cell lines. However, in the absence of IFN pre-treatment and despite an important amount of endogenous IFNs produced upon infection in HAE cells, SARS-CoV-2 replication was highly efficient in these cells. Similar data were obtained in Calu-3 cells, despite an earlier IFN response than in HAE cells. This suggested that IFNs were produced too late to efficiently prevent replication, as proposed in another study (Lei et al., 2020b) and/or did not efficiently induced ISG expression. In agreement with this, we observed that knocking-out genes essential for type I and III responses (e.g. IRF9 or JAK1) had not beneficial impact on replication in model lung cell lines, contrary to what was reported in intestinal model cell lines (Stanifer et al., 2020). Moreover, adding high amounts of exogenous type I IFN had no real impact on replication when IFN was added post-infection (even as early as 8 h post-infection, when a high MOI was used), supporting the idea that the timing of IFN exposure is key to control replication. In line with this, SARS-CoV-2 has been shown to efficiently dampen IFN responses and ISG induction through several mechanisms (Konno et al., 2020; Lei et al., 2020a; Miorin et al., 2020; Sa Ribero et al., 2020; Xia et al., 2020). Indeed, nsp1, nsp6, nsp13, ORF3a, M, ORF7a and ORF7b inhibit STAT1/2 phosphorylation and STAT1 nuclear translocation is inhibited by ORF6. Nonetheless, numerous clinical trials are currently evaluating the impact of IFN therapy on COVID-19 patients and should shed light on whether exogenous IFN could be useful in this context. Of note, a substantial proportion of patients with severe diseases may well be unresponsive to such treatments, due to the presence of anti-IFN autoantibodies or inborn mutations in genes belonging to the IFN pathway, such as *IFNAR1* (Bastard et al., 2020; Zhang et al., 2020). This highlights the importance to identify the IFN-induced antiviral effectors, which are so potently active against SARS-CoV-2, in order to potentially guide future, targeted therapeutic interventions.

## Materials and Methods

### Plasmids and constructs

The pRRL.sin.cPPT.SFFV/IRES-puro.WPRE lentiviral vector has been described (Doyle et al., 2018). Human ACE2 (NM_021804) and TMPRSS2 variant 1 (herein called TMPRSS2v1; NM_001135099) were amplified using the SuperScript® III One-Step RT-PCR System with Platinum® Taq (Invitrogen) from 500 ng RNA obtained from 293T cells and Caco-2 cells, using primers 5’-AATTAATTTAGCGGCCGCATGTCAAGCTCTTCCTGGCTCC-3’ and 5’-AATTAATTTACTCGAGCTAAAAGGAGGTCTGAACATCATCAGTG-3 and 5’-AATTAATTTAGCGGCCGCATGCCCCCTGCCCCGCC-3’ and 5’-AATTAATTTACTCGAGTTAGCCGTCTGCCCTCATTTGTC-3’, respectively, and digested by NotI and XhoI. Human ACE2 was inserted into NotI-XhoI-digested pRRL.sin.cPPT.SFFV/IRES-puro.WPRE to generate pRRL.sin.cPPT.SFFV/ACE2.IRES-puro.WPRE (Addgene 145839). The IRES-puromycinR cassette was removed by XhoI-SalI digestion and either replaced by nothing, to generate pRRL.sin.cPPT.SFFV/ACE2.WPRE (Addgene 145842), or by an IRES-neomycinR cassette, to generate pRRL.sin.cPPT.SFFV/ACE2.IRES-neo.WPRE (Addgene 145840), or by an IRES-hygromycin R cassette, to generate pRRL.sin.cPPT.SFFV/ACE2.IRES-hygro.WPRE (Addgene 145841), respectively. These cassettes were obtained by overlapping PCR using primers 5’-AATTAATTCTCGAGGTTAACGAATTCCGCCC-3’ and 5’-GTTCAATCATGGTTGTGGCCATATTATCATCGTGTTTTTC-3’ and 5’-ATATGGCCACAACCATGATTGAACAAGATGGATTGCACGC-3’ and 5’-TATATATTAGTCGACTCAGAAGAACTCGTCAAGAAGGCGATAG-3’ on the ECMV IRES sequence and the neomycin resistance gene (amplified using pRRL.sin.cPPT.SFFV/IRES-puro.WPRE and pcDNA3.1+, respectively) and using primers 5’-AATTAATTCTCGAGGTTAACGAATTCCGCCC-3’ and AGGCTTTTTCATGGTTGTGGCCATATTATCATCGTGTTTTTC-3’ and 5’-ATATGGCCACAACCATGAAAAAGCCTGAACTCACCGC-3’ and 5’-TTAATTAATTGTCGACCTATTCCTTTGCCCTCGGACGAGTG-3’ on the ECMV IRES sequence and the hygromycin resistance gene (amplified using pAHM (Goujon et al., 2013)), respectively. Human TMPRSS2 was cloned into NotI-XhoI-digested pRRL.sin.cPPT.SFFV/IRES-neo.WPRE to generate pRRL.sin.cPPT.SFFV/TMPRSS2v1.IRES-neo.WPRE (Addgene 145843). Of note, a mutation (G8V) is present on the cloned CDS of TMPRSS2v1 but this does not seem to impact functionality (not shown).

The pLX_311-Cas9 and LentiGuide-Puro vectors were gifts from John Doench and Feng Zhang, respectively (Doench et al., 2014; Sanjana et al., 2014) (Addgene 96924 and 52963) and we have described before the LentiGuide-Neo, LentiGuide-Neo-CTRLg1 and g2 (Doyle et al., 2018) (Addgene 139449, 139450, 139451). Guide RNA coding oligonucleotides were annealed and ligated into BsmBI-digested LentiGuide-Neo vector, as described (Addgene). The gRNA coding sequences used were as follow: gRIG-I 5’-GGGTCTTCCGGATATAATCC, gMDA-5 5’-TGGTTGGACTCGGGAATTCG; gMAVS 5’-AGGTGGCCCGCAGTCGATCC; gIRF9 5’-CAGCAACTGATACACCTTGT; and gJAK1 5’-TCTCGTCATACAGGGCAAAG.

### Cell lines

Human 293T, A549, Caco-2 and Calu-3, HEK-Blue™ IFN-α/β and IFN-γ cells, dog MDCK cells simian Vero E6 cells were maintained in complete Dulbecco’s modified Eagle medium (DMEM) (Gibco) supplemented with 10% foetal bovine serum and penicillin/streptomycin. HEK-Blue™ IFN-α/β and IFN-γ cells were cultured with 100 μg/ml zeocin and 30 μg/ml Blasticidin, or 100 μg/ml zeocin, 30 μg/ml Blasticidin and 1 μg/ml puromycin, respectively. Caco-2 and Calu-3 cells were obtained from American Type Culture Collection (ATCC); HEK-Blue™ IFN-α/β and IFN-γ cells were obtained from InvivoGen; 293T, MDCK, A549, Vero E6 cells were gifts from Michael Malim’s lab, Wendy Barclay’s lab, and from the CEMIPAI facility, respectively. A549 and Caco-2 cells stably expressing ACE2 and TMPRSS2 were generated by transduction with either RRL.sin.cPPT.SFFV/IRES-puro.WPRE, RRL.sin.cPPT.SFFV/IRES-neo.WPRE, RRL.sin.cPPT.SFFV/IRES-hygro.WPRE or RRL.sin.cPPT.SFFV.WPRE containing-vectors (cDNA as indicated) and were maintained under 1 μg/ml puromycin, and/or 1 mg/ml G418, 50 μg/ml hygromycinB selection.

For CRISPR-Cas9-mediated gene disruption, A549-ACE2 and Calu-3 cells stably expressing Cas9 were first generated by transduction with LX_311-Cas9 followed by blasticidin selection at 10 μg/ml. Cas9 activity was checked using the XPR_047 assay (a gift from David Root, Addgene 107145) and was 79.5% and >83.4%, respectively, for Cas9-expressing A549-ACE2 and Calu-3 cells. The cells were then transduced with guide RNA expressing LentiGuide-Puro and Lentiguide-Neo vectors (as indicated) and selected with antibiotics for at least 10 days.

Air-liquide cultures of primary Human Airway Epithelial (HAE) cells of nasal, tracheal and bronchial origins from healthy donors were obtained from Epithélix (MucilAir™) and cultured with MucilAir™ media (Epithélix). The apical side of the HAE cells was washed when necessary and 1 day prior to IFN exposure, following the manufacturer’s instructions.

When indicated, universal type I IFN (PBL Interferon source) was added at the indicated concentration (e.g. 1,000 U/ml) for 16-24 hr prior to virus infection. For HAE cells, IFN was added both in the basal media and on the apical side of the cells (diluted in 20 μl of MucilAir media; of note, 20 μl of media without IFN was added to the control cells in parallel).

### Lentiviral production and infection

Lentiviral vector stocks were obtained by polyethylenimine (PEI; for LentiGuides) or Lipofectamine 3000 (Thermo Scientific; for ACE2 and TMPRSS2 lentiviral vectors)-mediated multiple transfection of 293T cells in 6-well plates with vectors expressing Gag-Pol, the miniviral genome, the Env glycoprotein at a ratio of 1:1:0.5. The culture medium was changed 6 h post-transfection, and vector containing supernatants harvested 36 h later, filtered and used directly or stored at −80**°**C.

### SARS-CoV-2 production and infection

The BetaCoV/France/IDF0372/2020 isolate was supplied by Pr. Sylvie van der Werf and the National Reference Centre for Respiratory Viruses hosted by Institut Pasteur (Paris, France). The patient sample from which strain BetaCoV/France/IDF0372/2020 was isolated was provided by Dr. X. Lescure and Pr. Y. Yazdanpanah from the Bichat Hospital, Paris, France. The BetaCoV/France/IDF0372/2020 was amplified in Vero E6 cells (MOI 0,005) in serum-free media supplemented with 0,1 μg/ml L-1-p-Tosylamino-2-phenylethyl chloromethylketone (TPCK)-treated trypsin (Sigma–Aldrich). The supernatant was harvested at 72 h post infection when cytopathic effects were observed (with around 50% cell death), cell debris were removed by centrifugation, and aliquots frozen down at −80°C. Viral supernatants were titrated by plaque assays in Vero E6 cells. Typical titers were 3-5.10^6^ plaque forming units (PFU)/ml.

Simian Vero E6 and human cell infections were performed at the indicated multiplicity of infection (MOI; as calculated from titers in Vero E6 cells) in serum-free DMEM and 5% serum-containing DMEM, respectively. The viral input was left for the duration of the experiment (unless specified otherwise). The viral supernatants were frozen down at −80°C prior to RNA extraction and quantification and/or titration by plaque assays.

HAE cells were incubated for 2h with SARS-CoV-2 diluted in 50 μl of PBS1X added to the apical side. The viral input was then removed and the cells washed with 100 μl PBS1X. To collect the progeny viruses at 24 h, 48 h, and 72 h post infection, 50 μl of PBS1X was added to the apical side of the cells and collected after a 20 min incubation at 37°C. The viral supernatants were frozen down at −80°C prior to titration by plaque assays on Vero E6 cells. The cells were lysed in RLT buffer (Qiagen) followed by RNA extraction at 72 h post-infection.

### Quantification of mRNA expression

3-5 × 10^5^ cells with or without treatment with IFNα and SARS-COV-2 infection were harvested and total RNA was extracted using the RNeasy kit (Qiagen) employing on-column DNase treatment, according to the manufacturer’s instructions. 140 μl of supernatants from infected cells were subjected to RNA extraction using the QIAamp Viral RNA Mini Kit (Qiagen), according to the manufacturer’s instructions. 125 ng cellular RNA or 1,2 μl viral RNA were used to generate cDNAs. The cDNAs were analysed by qPCR using published RdRp primers and probe (Corman et al., 2020), as follow: RdRp_for 5’-GTGARATGGTCATGTGTGGCGG-3’, RdRp_rev 5’-CAAATGTTAAAAACACTATTAGCATA-3’ RdRp_probe 5’-FAM-CAGGTGGAACCTCATCAGGAGATGC-TAMRA-3’) and/or TaqMan gene expression assays (Applied Biosystems) for *ACTB* (Hs99999903_m1), *GAPDH* (Hs99999905_m1), *ISG15* (Hs01921425_s1), *OAS1* (Hs00973637_m1), *IFITM3* (Hs03057129_s1), *MX1* (Hs00182073_m1), *IFNB1* (Hs01077958_s1), *IFNL1* (Hs00601677_g1), *IFNL2* (Hs00820125_g1). qPCR reactions were performed in triplicate, in universal PCR master mix using 900 nM of each primer and 250 nM probe or the indicated Taqmans. After 10 min at 95°C, reactions were cycled through 15 s at 95°C followed by 1 min at 60°C for 40 repeats. Triplicate reactions were run according to the manufacturer’s instructions using a ViiA7 Real Time PCR system (ThermoFisher Scientific). For ISG expression, *GAPDH* and/or *ACTB* mRNA expression was used to normalize samples. pRdRp (which contains an RdRp fragment amplified from SARS-CoV-2 infected cell RNAs using primers RdRp_for and RdRp_rev and cloned into pPCR-Blunt II-TOPO) was diluted in 20 ng/ml salmon sperm DNA to generate a standard curve to calculate relative cDNA copy numbers and confirm the assay linearity (detection limit: 10 molecules of RdRp per reaction).

### RT^2^ profiler

The RT^2^ First Strand kit (Qiagen) was used for the synthesis of the complementary DNA strand using 400 μg of total RNA from samples extracted using the RNeasy kit (Qiagen) employing on-column DNase treatment. RT^2^ Profiler™ PCR Array Human Antiviral Response (PAHS-122Z) was used in the present study, according to manufacturer’s instructions. The ViiA7 Real Time PCR system (ThermoFisher Scientific) was used to amplify the DNA with a thermal cycling of 95 °C for 10 min followed by 40 cycles of 15 s at 95°C and 60 s at 60°C. Five housekeeping genes (β-actin (ACTB), β-2-microglobulin (B2M), glyceraldehyde-3-phosphate dehydrogenase (GAPDH), hypoxanthine phosphoribosyltransferase1 (HPRT1) and ribosomal protein, large, P0 (HPLP0) were used as internal controls. The average of the Ct values from these 5 controls was used to normalize gene expression. Changes in mRNA expression between the non-infected and the infected conditions were analysed using the ΔΔCt method.

### Quantification of secreted Cytokines

The concentration of 13 secreted cytokines was measured in the supernatants and basal media of infected Calu-3 and HAE cells, respectively, at the indicated conditions, using LEGENDplex bead-based immunoassays (BioLegend, human anti-virus response panel), according to the manufacturer’s recommendations. Samples were analysed on a BD Canto II flow cytometer using the Diva software (BD Biosciences, San Jose, CA). BioLegend’s LEGENDplex Data Analysis Software was used to analyse data.

### Immunofluorescence and microscopy

HAE cells were pre-treated or not with IFN for 20 h and infected with SARS-CoV2, as described above, for 48 h. Cells were fixed with PBS1X containing 4% paraformaldehyde (EM Sciences) for 15 min, permeabilized with 0.5% Triton X-100 for 15 min, and blocked/quenched in buffer NGB (50 mM NH_4_Cl, 1% goat serum, 1% bovine serum albumin) for 1 h. An overnight incubation at 4°C with mab J2 (Scicons) followed by incubation in a secondary anti-mouse antibody conjugated to Alexa Fluor 546 and in Alexa Fluor 488 Phalloidin (Thermofisher Scientific) for 2h at RT were used to visualise dsRNA and F-actin, respectively. The transwell membranes were removed from the inserts and mounted between slides and coverslips using ProLong™ Gold Antifade Mountant (Thermofisher Scientific). Images were acquired with a LSM880 confocal microscope paired with an Airyscan module (ZEISS) with a 63x lens. Post-processing of RAW Airyscan images was performed using the Zen Black software.

### HEK-Blue™ IFN-α/β and IFN-γ assays

HEK-Blue™ IFN-α/β cells and HEK-Blue™ IFN-γ (InvivoGen) were plated at 30,000 cells per well in a 96-well plate. The following day, media from infected cells (or control cells) was added and a standard curve was generated in parallel by serial dilutions of type I or type III IFNs in complete DMEM. After 20-24 h incubation, 30 μl of HEK-Blue™ IFN-α/β supernatants was added to 120 uL of Quanti-blue™ substrate (InvivoGen) and incubated at 37°C for 15 min. Absorbance was measured at 620nm using an Envision plate reader (Perkin-Elmer). The standard curves were used to provide semi-quantitative analyses of the IFN concentrations produced by the infected cells.

### Spike intracellular staining and flow cytometry analysis

Infected cells were harvested at the indicated time points post-infection and fixed for 30 min in PBS1X-4% PFA. Cells were washed once in PBS1X and twice in BD Perm-Wash buffer and permeabilized for 15 min at RT in BD Perm-Wash buffer. Cells were incubated on ice for 30-45 min in FACS buffer (PBS1X-5% FCS) containing a 1/250 dilution of Alexa 488-conjugated anti-Spike antibody (GTX632604 conjugated using the Zenon Alexa Fluor 488 Mouse IgG Labeling Kit, ThermoFisher) and washed 4 times in FACS buffer. Flow cytometry was performed using the NovoCyte™ (ACEA Biosciences Inc.).

### Immunoblot analysis

Cells were lysed in lysis buffer (10 mM TRIS 1M pH7.6, NaCl 150 mM, Triton X100 1%, EDTA 1 mM, deoxycholate 0,1%) supplemented with sample buffer (50 mM Tris-HCl pH 6.8, 2% SDS, 5% glycerol, 100 mM DTT, 0.02% bromphenol blue), resolved by SDS-PAGE and analysed by immunoblotting using primary antibodies against SARS-CoV Nucleocapsid (Bio-Techne NB100-56683), SARS-CoV Spike (GeneTex GTX632604), Actin (Sigma-Aldrich A1978), IFITM3 (Proteintech 11714-1-AP), MX1 (ThermoFisher Scientific PA5-22101), RIG-I (Covalab mab10110), MDA-5 (Ozyme D74E4), and MAVS (ProteinTech 14341-1-AP), followed by secondary horseradish peroxidase-conjugated anti-mouse or anti-rabbit immunoglobulin antibodies and chemiluminescence Clarity or Clarity max substrate (Bio-Rad). A Bio-Rad ChemiDoc imager was used.

## Data availability

The datasets generated during and/or analysed during the current study are available from the corresponding authors on reasonable request.

## Requests for materials

Requests for material should be addressed to Caroline Goujon or Olivier Moncorgé at the corresponding address above, or to Addgene for the plasmids with an Addgene number.

## Acknowledgements

We wish to thank Raphaël Gaudin, Laurence Briant, Sylvie Van der Werf, Eric Bernard, Emma Partiot, Aurélie Lieuvin, Hélène Bauby, Mélanie Wencker for the generous provision of reagents, protocols and/or for helpful discussions. We are immensely grateful to Christine Chable-Bessia for setting up excellent working conditions for SARS-CoV-2 handling at the CEMIPAI BSL-3 facility.

This work was supported by the European Research Council (ERC) under the European Union’s Horizon 2020 research and innovation programme (grant agreements n°759226 and n°899835) (to CG), the Institut National de la Santé et de la Recherche Médicale (INSERM) (to CG), the ATIP-Avenir programme (to CG) and institutional funds from the Centre National de la Recherche Scientifique (CNRS) and Institut des Sciences Biologiques du CNRS (INSB) (to CG and OM), the French National Research Agency ANR under the “Investissements d’avenir” programme with the reference ANR-16-IDEX-0006 through Montpellier University MUSE program (to CG), the Labex EpiGenMed, an Investissements d’avenir program (reference ANR-10-LABX-12-01) (to SN), the Région Occitanie and the Agence Nationale de la Recherche (ANR Flash COVID-19, reference ANR-20-COVI-0099) (to SN), 3-year PhD studentships from the Ministry of Higher Education and Research (to AR, to BB and to JM), a 4th year PhD funding from the Fondation pour la Recherche Médicale (to BB), a 2-year Sidaction post-doctoral fellowship (to R.P.) and a 2-year post-doctoral grant from the Agence Nationale de la Recherche sur le SIDA et les Hépatites virales, ANRS (to GM). We acknowledge the imaging facility MRI, member of the national infrastructure France-BioImaging supported by the French National Research Agency (ANR-10-INBS-04) and the CEMIPAI BSL-3 facility.

## Author contribution statement

A.R., O.M., and C.G. conceived and designed the experiments; A.R., A.L.C.V., M.T., O.M., and C.G. performed the main experiments; G.M. and S.N. performed the LegendPlex experiments and analyses; B.B. designed the molecular cloning strategies; J.M. did the immunofluorescence and microscopy analyses; R.P., and M.A. provided technical help; A.R., O.M. and C.G. wrote the manuscript with input from all authors.

## Supplementary Information

**Figure S1.**
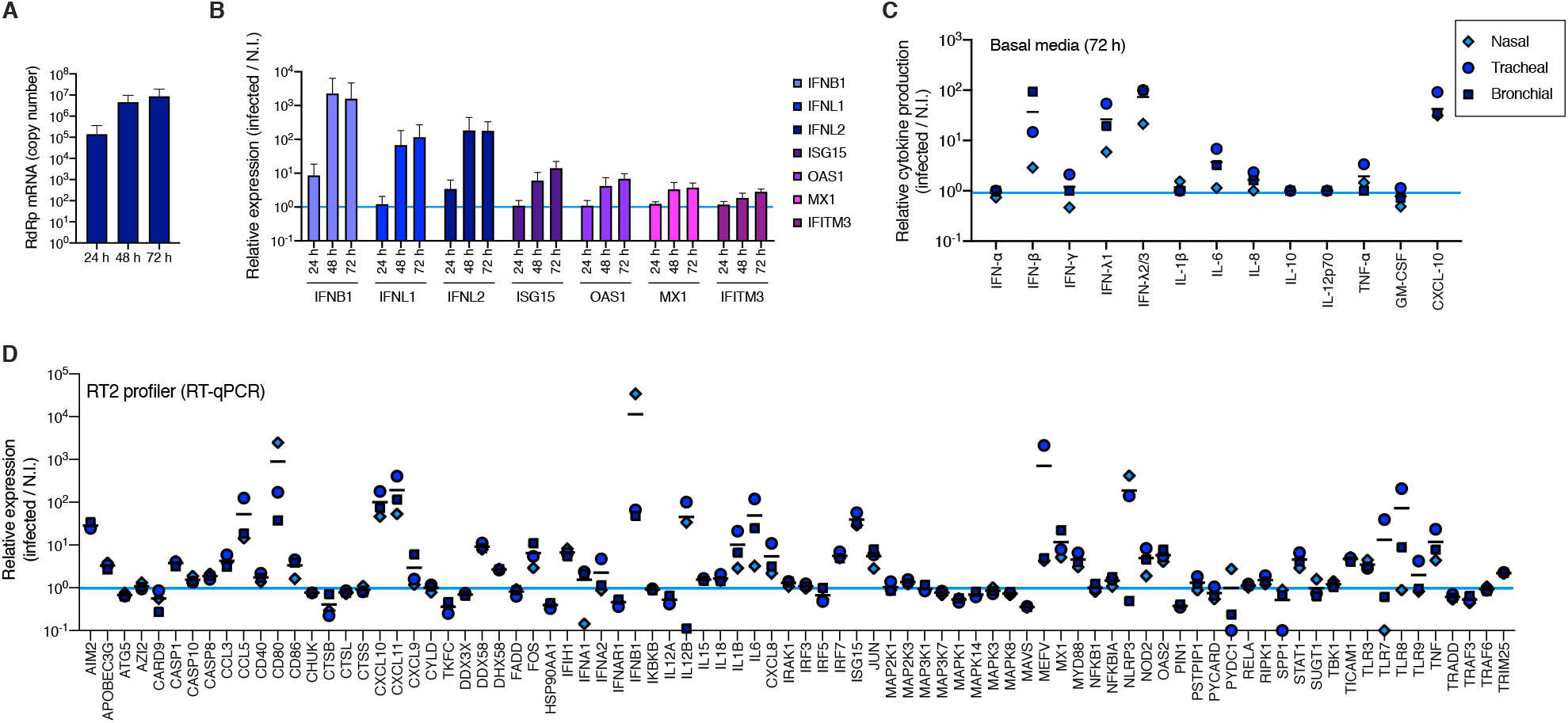
Primary human airway epithelial host cell responses to SARS-CoV-2 infection. **A**. Human HAE cells of nasal, tracheal and bronchial origins were mock infected (N.I.) or incubated with SARS-CoV-2 on the apical side at MOI 0,01 for 2 h as in Figure 1. Cells were harvested at the indicated time points and lysed for RNA extraction and RT-qPCR analysis using RdRp primers and probe. **B**. Differential gene expression was measured in RNAs from A. using the indicated taqmans, and data were normalized to both ActinB and GAPDH. **C**. Data from Human Anti-Virus Response Panel LEGENDplex™ as performed in Figure 1A with supernatants from cells of nasal, tracheal and bronchial origins. **D**. Data from Antiviral Response RT^2^ profiler PCR array analysis as in Figure 1D for RNAs from cells of nasal, tracheal and bronchial origins. The light blue line (sets at 1) indicates no change in cytokine production or in gene expression (C and D). The mean of 6 (A and B) or 3 (C and D) independent experiments is shown (apart for the 24 h time point in A and B, n=3), with error bars representing one standard deviation (s.d.) from the mean (A and B).

**Figure S2.**
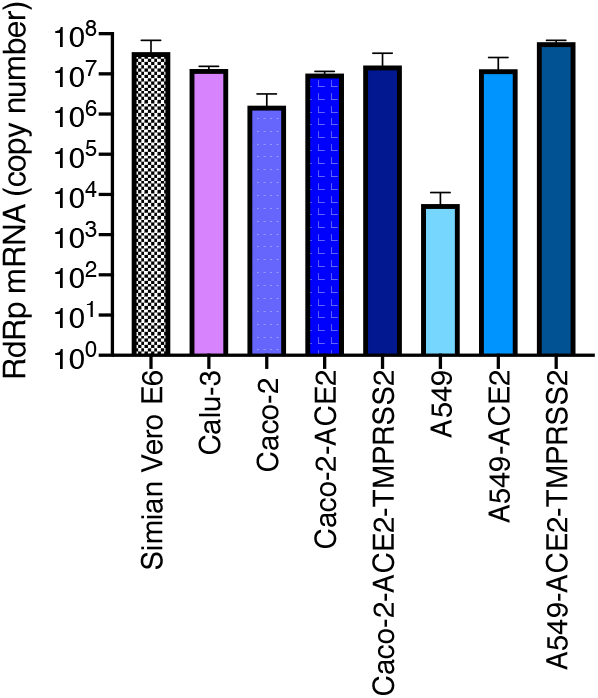
Replication of SARS-CoV-2 in genetically modified human cell lines. Caco-2 and A549 cells were transduced or not with lentiviral vectors to stably overexpress ACE2, or ACE2 and TMPRSS2. The indicated (unmodified and modified) cell lines were infected with SARS-CoV-2 at MOI 0,05 and lysed 48 h later for RNA extraction and RdRp RT-qPCR analysis. A representative experiment (with technical triplicates) is shown.

**Figure S3:**
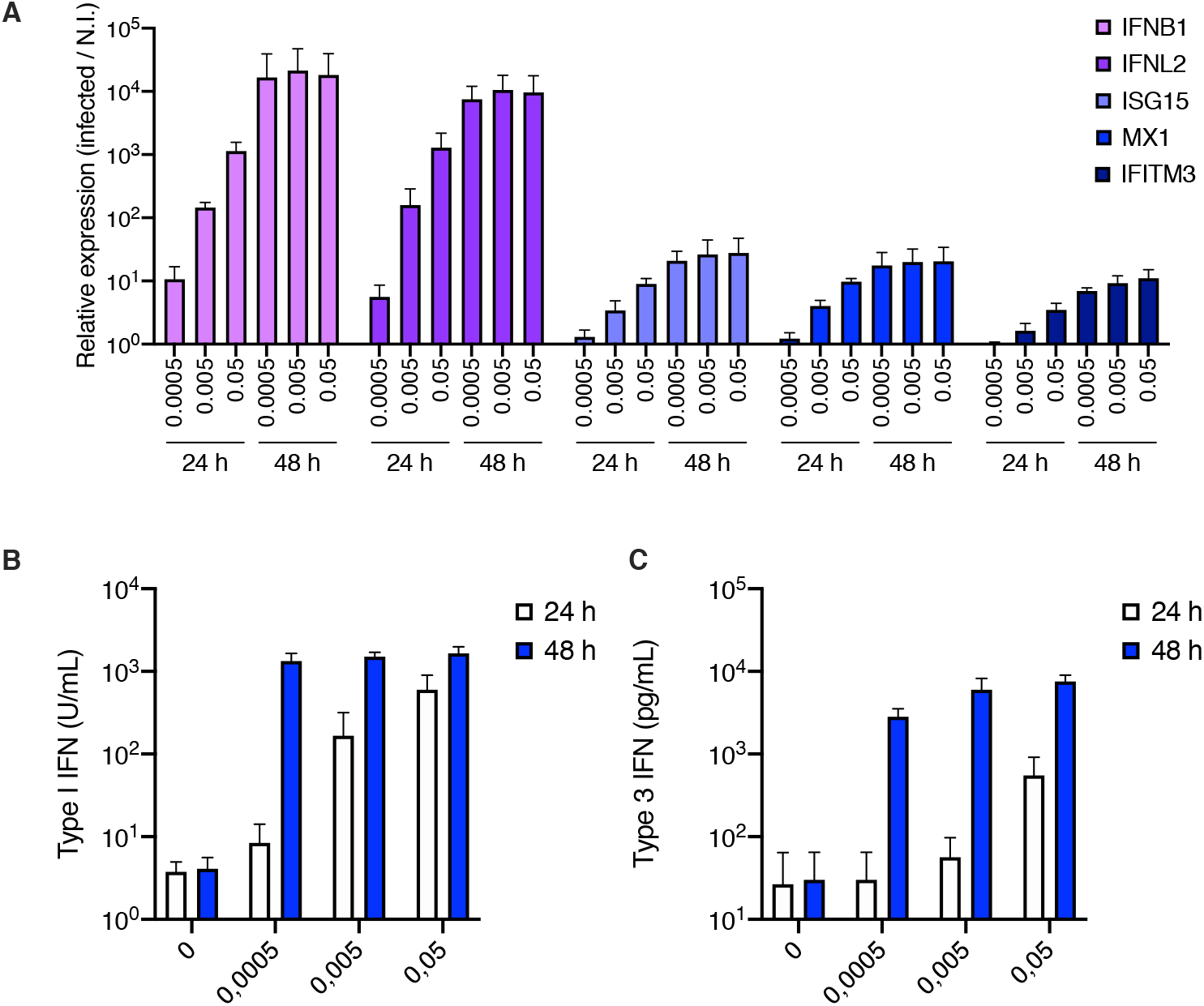
Type I and III interferon induction in Calu-3 cells infected by SARS-CoV-2. **A**. Human Calu-3 cells were mock infected or incubated with SARS-CoV-2 at the indicated MOIs. Cells were harvested at 24 h or 48 h and lysed for RNA extraction. Relative expression levels of the indicated IFN genes and ISGs were analysed by RT-qPCR analysis using both ActinB and GAPDH for normalization. **B**. The cell supernatants from (A) were harvested at the indicated time points and type I and type III IFN concentrations were measured using HEK-Blue™ IFN-α/β and IFN-γ reporter cells, respectively. The mean of 3 to 4 (A) or 3 (B and C) independent experiments is shown, with error bars representing one s.d. from the mean.

**Figure S4:**
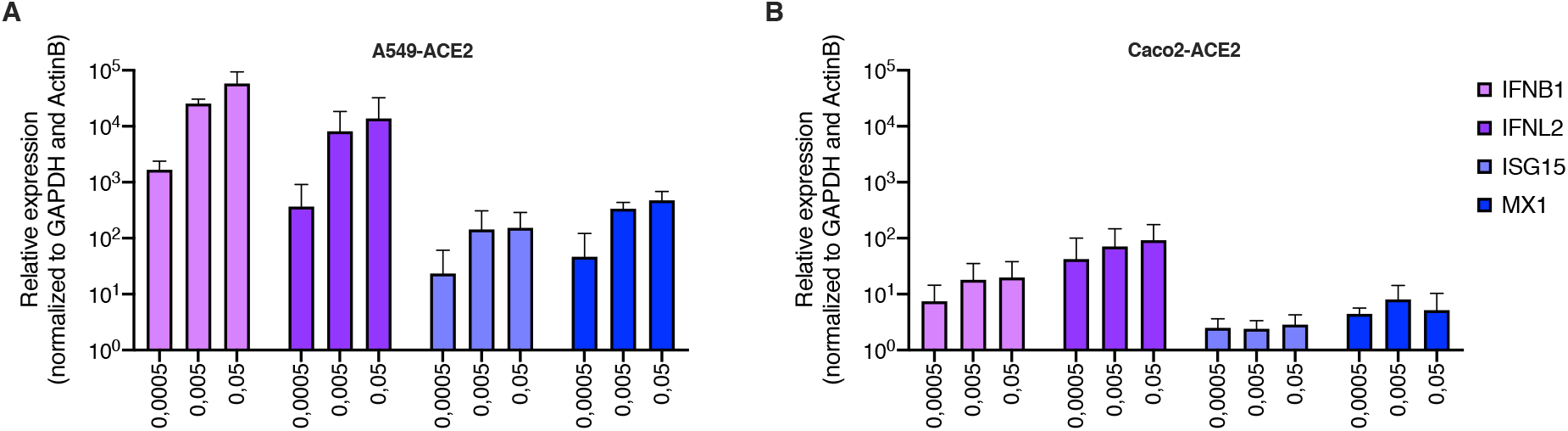
IFN gene and ISG induction in A549-ACE2 and Caco-2-ACE2 cells infected by SARS-CoV-2. Human A549-ACE2 (**A**) and Caco-2-ACE2 (**B**) cells were non-infected or incubated with SARS-CoV-2 at the indicated MOIs. Cells were harvested at 48 h and 24 h, respectively (i.e. the time points at which replication reached similar levels without a major impact on cell survival, not shown), and lysed for RNA extraction. Relative expression levels of the indicated IFN genes and ISGs were analysed by RT-qPCR analysis using both ActinB and GAPDH for normalization. The mean of 3 independent experiments is shown, with error bars representing one s.d. from the mean.

**Figure S5.**
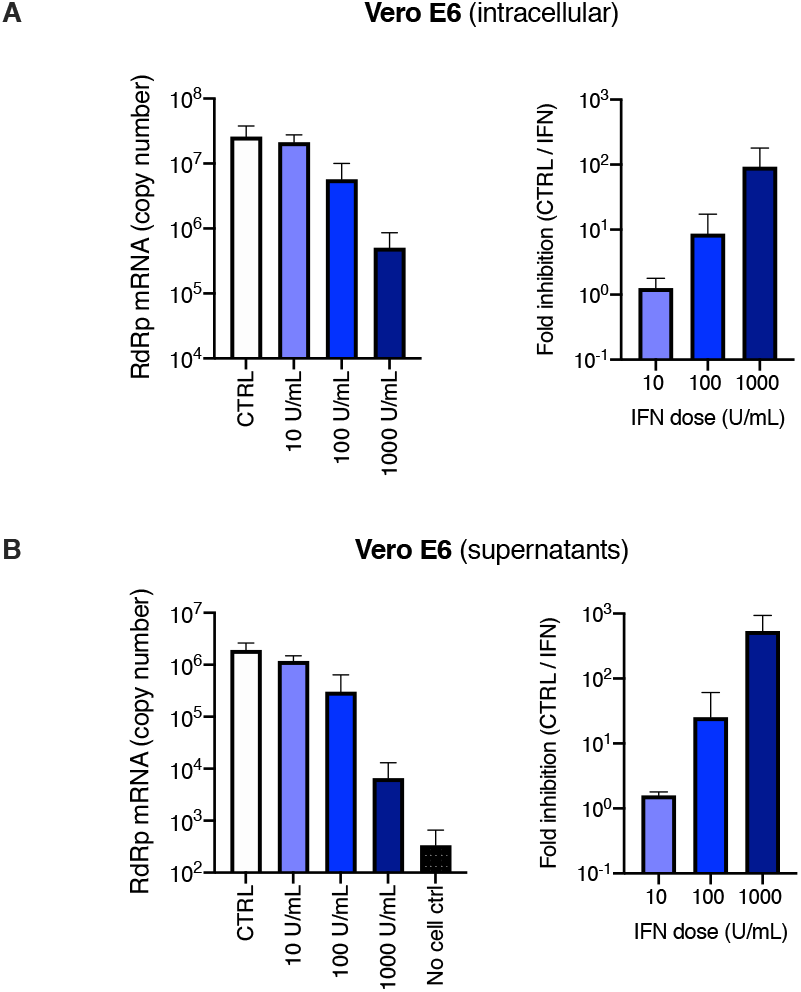
Inhibition of SARS-CoV-2 replication by type I IFN in Vero E6 cells. Vero E6 cells were pre-treated or not with increasing concentrations of type I IFN, as indicated, for 16 h prior to SARS-CoV-2 infection at MOI 0,0005. 72 h later, the cells were lysed and the supernatants collected, the RNAs were extracted and viral replication was monitored in cells (**A**, left panel) and viral production in the supernatants (**B**, left panel) by RdRp RT-qPCR. Right panels. The fold inhibition by IFN is shown (**A** and **B**, right panels). The mean of 3 independent experiments is shown, with error bars representing one s.d. from the mean.

**Figure S6.**
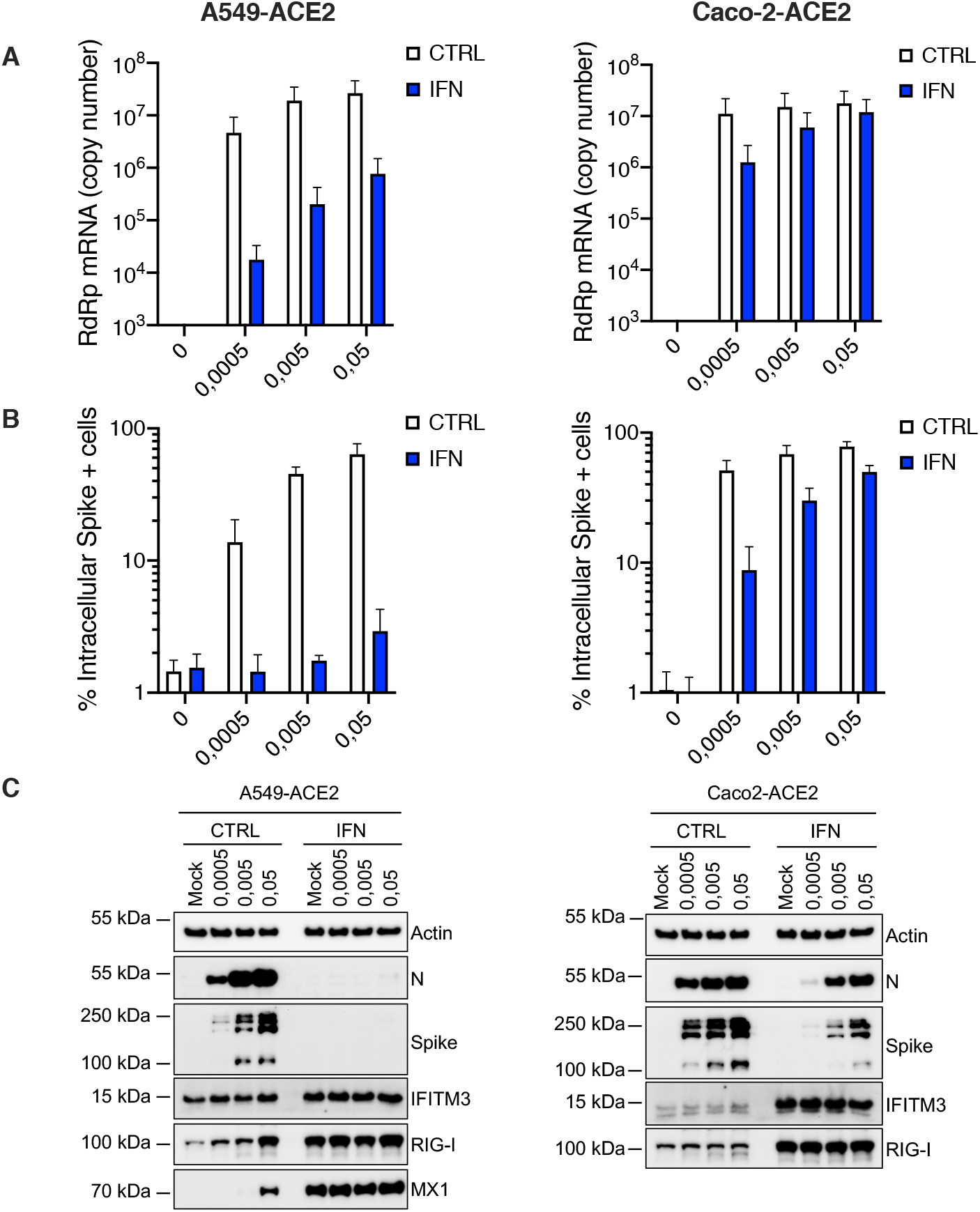
Inhibition of SARS-CoV-2 replication by type I IFN pre-treatment in A549-ACE2 and Caco-2-ACE2 cell lines. Human A549-ACE2 (left) and Caco-2-ACE2 (right) cells were pre-treated or not with IFN for 16-20 h, the media was replaced and the cells were mock-infected (N.I.) or incubated with SARS-CoV-2 at the indicated MOIs. Cells were harvested at 48 h (A549-ACE2) or 24 h (Caco-2-ACE2). **A**. The cells were lysed for RNA extraction and RT-qPCR analysis using RdRp primers and probe. **B**. The cells were fixed with PFA at 24 h post-infection, permeabilized and stained with an anti-Spike antibody conjugated to an Alexa fluorochrome. The percentage of Spike + cells was scored by flow cytometry. **C**. The cells were lysed for Immunoblot analysis of SARS-CoV-2 Nucleoprotein (N) and Spike, IFITM3, RIG-I and MX1 ISG expression levels, Actin serving as a loading control. Of note, MX1 was not detected in Caco-2-ACE2 cell lysates. A representative immunoblot is shown. The mean of 3 independent experiments is shown (A and B), with error bars representing one standard deviation (s.d.) from the mean.

**Figure S7.**
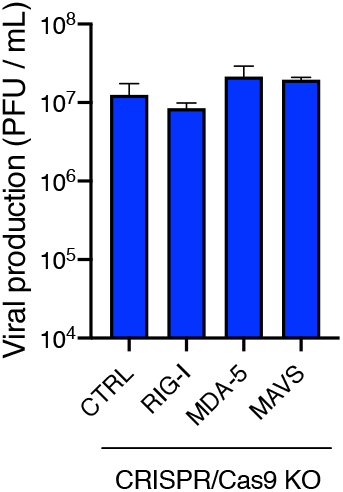
MDA-5 and MAVS knockout do not impact viral production in Calu-3 cells. CTRL, RIG-I, MDA-5 and MAVS Calu-3 knockout cells were infected with SARS-CoV-2 at MOI 0,05 (as in Figure 3) and viral production was measured 48 h later by plaque assays on Vero E6 cells. The mean of 2 independent experiments is shown, with error bars representing one s.d. from the mean.

**Figure S8.**
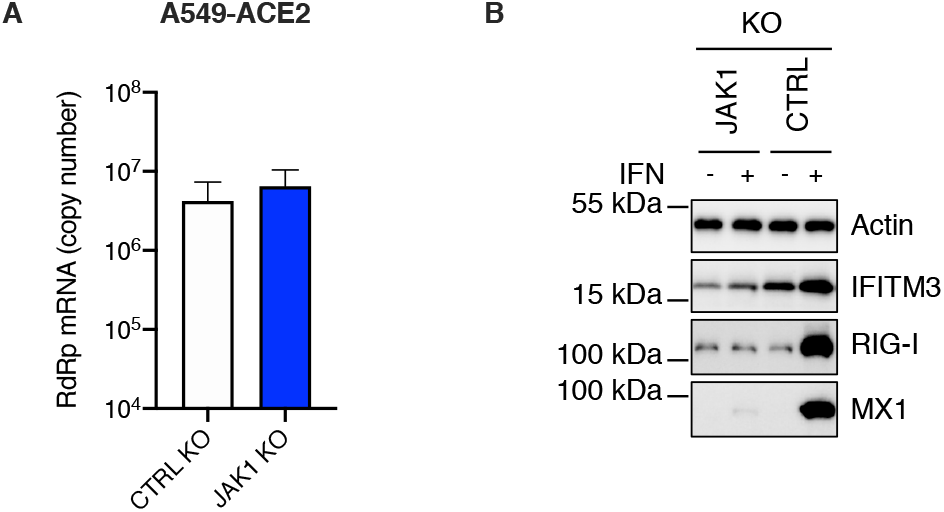
The induction of interferons does not contribute to controlling SARS-CoV-2 replication in A549-ACE2 cells. **A**. CTRL and JAK1 A549-ACE2 knockout cells were infected with SARS-CoV-2 at MOI 0,0005 and viral replication was measured 48 h later using RdRp RT-qPCR. The mean of 2 independent experiments is shown, with error bars representing one s.d. from the mean. **B**. CTRL and JAK1 knockout cells were pre-treated or not with IFN for 48 h, lysed and the expression levels of IFITM3, RIG-I and MX1 were analysed by immunoblot, Actin serving as a loading control. A representative immunoblot is shown.

**Figure S9.**
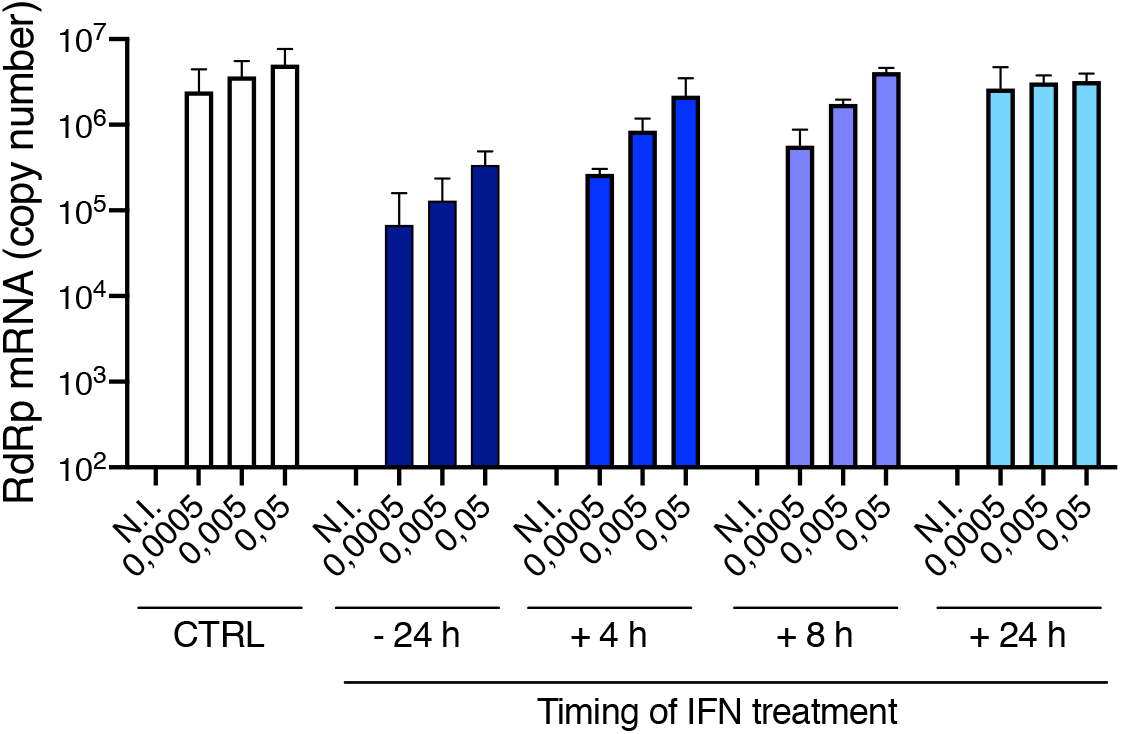
Type I interferons do not help controlling SARS-CoV-2 replication when added post-infection. Calu-3 cells were infected with SARS-CoV-2 at MOI 0,0005, 0,005 and 0,05 after a 24h pre-treatment with IFN or not, or were subsequently treated with IFN at 4 h, 8 h or 24 h post infection. Viral replication was measured 48 h post infection using RdRp RT-qPCR. The mean of 3 independent experiments is shown, with error bars representing one s.d. from the mean.

